# Serum Amyloid A proteins reduce bone mass during mycobacterial infections

**DOI:** 10.1101/2022.10.24.513637

**Authors:** Ana Cordeiro Gomes, Daniela Monteiro Sousa, Tiago Carvalho Oliveira, Óscar Fonseca, Ricardo J. Pinto, Diogo Silvério, Ana Isabel Fernandes, Ana C. Moreira, Tânia Silva, Maria José Teles, Luísa Pereira, Margarida Saraiva, Meriem Lamghari, Maria Salomé Gomes

## Abstract

Osteopenia has been associated to several inflammatory conditions, including mycobacterial infections. How mycobacteria cause bone loss remains elusive, but direct bone infection may not be required. Using genetically engineered mice and morphometric, transcriptomic and functional analyses, we found that infection with *Mycobacterium avium* impacts bone turnover by decreasing bone formation and increasing bone resorption, in a IFNg- and TNFa-dependent manner. IFNg produced during infection enhanced macrophage TNFa secretion, which in turn increased the production of serum amyloid A (SAA) 3. *Saa3* expression was upregulated in the bone of both *M. avium*- and *Mycobacterium tuberculosis-infected* mice and SAA proteins were increased in the serum of patients with active tuberculosis. Furthermore, the increased SAA levels seen in active tuberculosis patients correlated with altered serum bone turnover markers. Additionally, human SAA proteins impaired bone matrix deposition and increased osteoclastogenesis in vitro. Overall, we report a novel crosstalk between the cytokine network operating in macrophages and bone homeostasis and disclose SAA proteins as potential biomarkers of bone loss during infection by mycobacteria.

## Introduction

A regulated balance between bone degradation and formation maintains an adequate bone mass, conferring the necessary rigidity and strength to the skeleton, to bear weight and avoid low impact fractures. Besides bone forming (osteoblasts and osteocytes) and bone degrading (osteoclasts) cells, bone remodeling is also influenced by immune mediators (Redlich and Smolen 2012, Oliveira et al. 2020). In support of this, loss of bone homeostasis during several inflammatory syndromes has been reported (Redlich and Smolen 2012), but the underlying mechanisms remain elusive. In this context, mycobacterial infections may present as good models to elucidate the molecular cross-talk between the immune response and bone homeostasis. The presence of mycobacteria in the bone has been traced back to thousands of years ago (Hogan et al. 2019). Mendelian susceptibility to mycobacterial disease is associated with a higher frequency of bone involvement (Tsumura et al. 2022). Furthermore, previous history of mycobacterial infection, even without bone involvement, increases the risk of osteopenia and osteoporosis (Yeh et al. 2016, Choi et al. 2017). Additionally, we and others have previously shown that mycobacterial infection skews the bone marrow hematopoietic development with important immune and hematological consequences (Baldridge et al. 2010, Matatall et al. 2014, Matatall et al. 2016, Oliveira et al. 2020, Gomes et al. 2021). Therefore, the infection of bone cells themselves or the immune modulation of the bone marrow niche during infection may disrupt bone mass (Oliveira et al. 2020).

In this study, we investigated the mechanisms underlying reduced bone mass during mycobacterial infection to shed light on how the immune system and bone homeostasis articulate. We combined morphometry, transcriptomic and functional in vitro and in vivo assays to study how infections with the non-tuberculous mycobacteria, *Mycobacterium avium*, impacted the bone. We established a mechanistic network centered in macrophages and linking IFNg, TNFa and serum amyloid A (SAA) proteins, which combined to dysregulate bone turnover. The production of SAA by macrophages was validated upon direct administration of TNFa to macrophages and during aerosol infection with *Mycobacterium tuberculosis*. Finally, we showed a systemic increase of SAA levels in patients with active tuberculosis, which positively correlated with altered bone turnover markers measured in the plasma. Our study advances our understanding on how the immune response to infection impacts bone loss and reveals potential clinical biomarkers for this. These findings are particularly important in tuberculosis disease, given the enormous number of cases in the world (WHO 2021), combined with an aging population more susceptible to bone disease.

## Results

### Mycobacterial infection reduces bone mass and mineral density due to both increased degradation and decreased formation

Given the body of evidence linking mycobacteria infections with bone involvement, we started by comprehensively addressing the alterations induced to the bone during *M. avium* infections. For that, C57BL/6J mice were infected with *M. avium* through the iv route (Gomes et al. 2019). Eight weeks later several parameters in the cortical and trabecular bone were quantified by micro-computerized tomography (microCT, **Fig. S1, A** to **C**). Even though the marrow volume (M.V.) and periosteal perimeter (Perio. P.) were increased in the cortical bone (**Fig. 1, A and B**), the cortical volume (C.V.), mineral density (C. B.M.D.) and thickness were reduced by infection (**Fig. 1, C** to **E**). Endocortical perimeter (Endo. P.) and mean polar moment of inertia (MPPI) were not altered (**Fig. 1, F** and **G**). Additionally, the trabecular bone volume (T.B.V.) was decreased (**Fig. 1H**) whereas the trabecular tissue volume (T.T.V.) was not altered (**Fig. 1I**), which resulted in a reduction of the ratio trabecular bone volume to tissue volume (%B.V./T.V., **Fig. 1J**). Also reduced in infected mice were the trabecular mineral density (T. B.M.D.), and the trabecular thickness and number (#T., **Fig. 1, K** to **M**), with the concomitant increase in the trabecular separation (T. S.; **Fig. 1N**).

**Figure 1.**
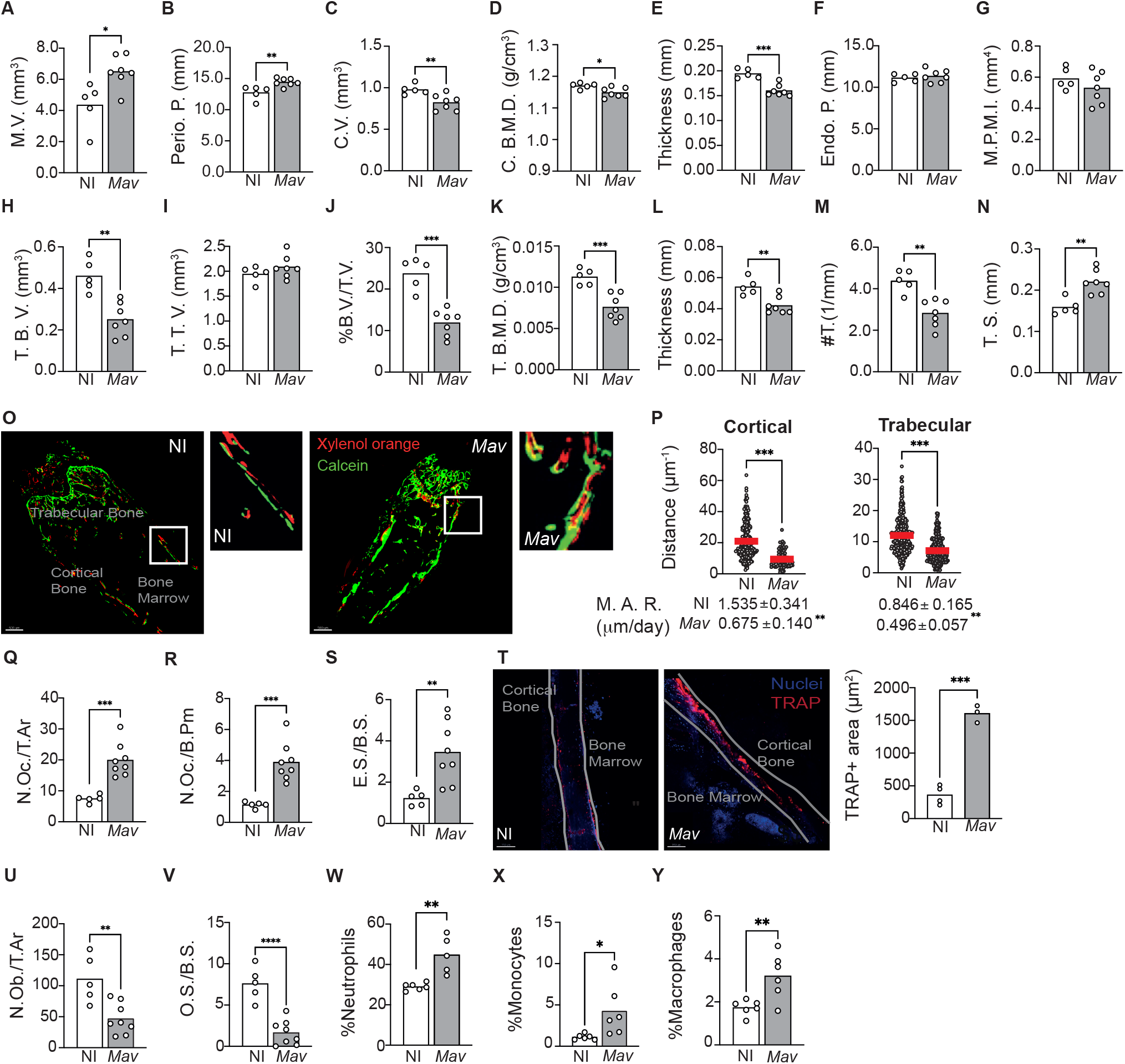
*Mycobacterium avium* infection alters bone turnover, reducing bone mass. **A-G)** Measurement of the marrow volume (M.V., **A**), periosteal perimeter (Perio. P., **B**), cortical volume (C.V., **C**), cortical bone mineral density (C. B. M. D., **D**), cortical thickness (**E**), endocortical perimeter (Endo. P., **C**), and polar mean moment of inertia (M. P. M. I., **G**) in *M. avium-infected* mice (*Mav* bars) compared to non-infected littermate controls (NI bars). **H-N**) Measurement of the trabecular bone volume (T.B.V., **H**), trabecular tissue volume (T.T.V., **I**), ratio of bone volume to tissue volume (%B.V./T.V., **J**), trabecular bone mineral bone density (T. B. M. D., **F**), trabecular thickness (**L**), trabecular number (#T, **M**) and separation (**N**) in *M. avium*-infected mice (*Mav* bars) and non-infected controls (NI bars).N=5-6 mice per experimental group; data representative of two independent experiments. Bars indicate average, and each circle depict each individual mouse analyzed. **O**) 20-μm thick sections of femoral bone labeled with xylenol orange (red) and calcein (green). Scale bar: 100μm. **P**) Distance between xylenol orange and calcein layers on the bone surface of non-infected (NI, black dots) and infected (*Mav*, white dots) mice. Bellow the graphs, the mineral apposition rate (M.A.R) is indicated. N=5-6 mice per experimental group. Red lines indicate average, and circles depict each measure. **Q-S**) Histomorphometry analysis of the long bones of *M. avium*-infected mice (*Mav* bars) compared with non-infected controls (NI bars). N. Oc/T.Ar, Osteoclast number per total area (**Q**), N.Oc./B.Pm., number of osteoclasts per bone perimeter (**R**), ES/BS, eroded surface per bone surface (**S**). N=6 mice per experimental group. Representative of two independent experiments. Bars indicate the average; circles depict each individual mouse. **T**) Quantification of the bone area expressing TRAP. TRAP-expressing bone surface were identified in the bones infected (*Mav*) and non-infected (NI) mice and the area of those regions was then measured. TRAP staining in red; nuclei in blue; white line shows the limits of the bone; NI, non-infected; *Mav*, infected for 8 weeks; Scale bar: 150μm. Bars represent the average of the experimental group; each dot represents each bone analyzed. **U-V)** Histomorphometry analysis of the long bones of *M. avium*-infected mice (*Mav* bars) compared with non-infected controls (NI bars). N.Ob/T.Ar, osteoblast number per total area (**U**), OS/BS, osteoid surface per bone surface (**V**). **W-Y**) Enumeration of neutrophils (Gr-1^hi^ CD115^−^; **W**), monocytes (Gr1^lo^ CD115^+^ F4/80^+^; **X**), and macrophages (Gr1^lo^ CD115^−^ F4/80^+^ CD169^+^; **Y**). N=6 mice per experimental group. Representative of three independent experiments. Bars indicate the average; circles depict each individual mouse. *, p<0.0,5, ** p<0.01, ***, p<0.001 by unpaired Student’s *t* test.

Given these alterations suggestive of bone loss during infection, a dynamic histomorphology analysis was performed (van Gaalen et al. 2010). Bone surface was fluorescently labeled in vivo by the injection of xylenol orange and calcein green at 6- and 8-weeks post-infection, respectively. Mice were sacrificed 3 days later, and 20-μm bone sections were analyzed by confocal microscopy. In non-infected mice, the two fluorochrome labels were detected in the bone and were visibly separated by non-labeled bone tissue. However, in the infected bones, even though both calcein and xylenol orange were detected, some regions were not labeled with xylenol orange, or the two layers of dyes overlapped (**Fig. 1O**). The distance between the xylenol and calcein layers in cortical and trabecular bone was measured and found to be significantly decreased in infected as compared to non-infected mice; concomitantly, the mineral appositional rate (M.A.R.) was decreased in infected mice (**Fig. 1P**).

Finally, histomorphology analysis of the bone revealed an increased number of osteoclasts per total area (N.Oc./T.Ar.) and per total bone perimeter (N.Oc./B.Pm., **Fig. 1, Q** and **R**), which led to an increased eroded bone surface per bone surface (E.S./B.S.; **Fig. 1S**). Together these findings indicate that bone loss is happening due to exacerbated osteoclast activity. To test this hypothesis, we measured the area of TRAP+ bone, which is indicative of osteoclast activity as TRAP is an enzyme secreted by osteoclasts during bone resorption. The area of TRAP+ bone surface was increased in infected mice (**Fig. 1T**). Furthermore, significantly higher levels of C-terminal telopeptides of Type I collagen (CTX-I) were detected in the serum of infected mice (**Fig. S1D**), an indicator of increased bone resorption.

In what regards the cellularity of the bone, we found that infection reduced the number of osteoblasts (N.Ob/T.Ar.) and osteoid volume per bone surface (O.S./B.S.) (**Fig. 1, U** and **V**), as well as the number of osteoblasts lining the bone surface (N.Ob./B.Pm.; **Fig. S1E**). The alterations in osteoblast lineage could also be detected at the LEPR^+^ mesenchymal stem cells (MSC) level, whose frequency was increased in the bone marrow of infected mice (**Fig. S1F**). LEPR^+^ MSC were described as the major source of osteoblasts and adipocytes in the adult bone marrow (Zhou et al. 2014) and nurse hematopoietic maintenance and differentiation (Cordeiro Gomes et al. 2016, Miao et al. 2022), reinforcing the close relationship between hematopoiesis and the bone (Tsumura et al. 2022).

Disseminated *M. avium* infection leads to the colonization of macrophages within the bone marrow parenchyma (Gomes et al. 2019) instead of a direct colonization of bone cells (**Fig. S2**), suggesting that the altered bone turnover and reduced bone mass that we now show during infection might be a consequence of the ongoing immune response. Previous reports have implied the bone marrow as a niche for mycobacteria (Beamer et al. 2014, Mayito et al. 2019), even during intranasal infection (Kager et al. 2014). Besides macrophages, hematopoietic (Tornack et al. 2017) and mesenchymal (Das et al. 2013) stem cells are colonized by mycobacteria in the bone marrow, which may explain the hematopoietic (Gomes et al. 2021) and bone dysfunctions. Of note, mycobacterial infections skew the differentiation of hematopoietic stem cells towards the myeloid lineage (Baldridge et al. 2010, Matatall et al. 2014, Matatall et al. 2016). Thus, we questioned whether alterations to the myeloid composition of the bone marrow were seen in our system. For this purpose, myeloid populations were identified as previously described (Chow et al. 2011). We found an increased frequency of neutrophils, monocytes and macrophages are present in the bone marrow of infected mice (**Fig. 1, W** to **Y**).

### IFNγ production during *M. avium* infection induces TNFa secretion leading to bone resorption

To gain insight into the molecular mechanisms behind reduced bone mass during mycobacterial infection, we performed targeted RNAseq of whole bone and compared the gene expression between non-infected and infected mice. We found that 1822 genes were statistical and differentially expressed (adjusted *P*-values of ≤ 0.05) in the bone of infected mice (**Fig. 2A**). Within the top upregulated differentially expressed genes (DEG) were several immune mediators, as *Cxcl9, Cxcl10* and *Nos2* (**Fig. 2A**). Furthermore, pathway-enrichment analysis revealed that the most upregulated (and statistically significant) pathways during *M. avium* infection were “response to IFNγ” and “IFNγ mediated signaling pathway” (**Fig. 2B**). As the hematological alterations driven by *M. avium* infections are known to be IFNγ-dependent (Baldridge et al. 2010, Matatall et al. 2016, Gomes et al. 2019), we next investigated whether this cytokine was also involved in bone loss during infection. For that, *Ifng^−/−^* mice were infected with *M. avium*. MicroCT analysis revealed that several measurements of cortical and trabecular bone found altered during infection of *Ifng^+/+^* (**Fig. 1**) were similar between infected and non-infected mice lacking *Ifng* (**Fig. 2, C** to **F**; and **Fig. S1, G** to **P)**. Thus, IFNγ is a central piece mediating bone loss during *M. avium* infection. Following IFNγ-related pathways, our RNA-Seq data highlighted an upregulation of pathways associated with TNF and NIK and NFkB signaling in the bone of infected mice (**Fig. 2B**). Since these pathways are important for osteoclast formation (Redlich and Smolen 2012, Abu-Amer 2013, Nevius et al. 2016, Amarasekara et al. 2018, Oliveira et al. 2020), we next measured the expression of genes encoding molecules involved in osteoclast differentiation. *M. avium* infection induced the up-regulation of *Csf1, Csf1r, Rank*, and *Acp5* in the bones of *Ifng^+/+^* but not of *Ifng^−/−^* mice (**Fig. 2**, **G** to **J)**. Overall, our data indicate that osteopenia during *M. avium* infection depends on IFNg production, which mediates the upregulation of genes related to osteoclast formation and activity. These observations agree with previous studies reporting that IFNg is an indirect promoter of osteoclastogenesis and bone resorption (Gao et al. 2007).

**Figure 2.**
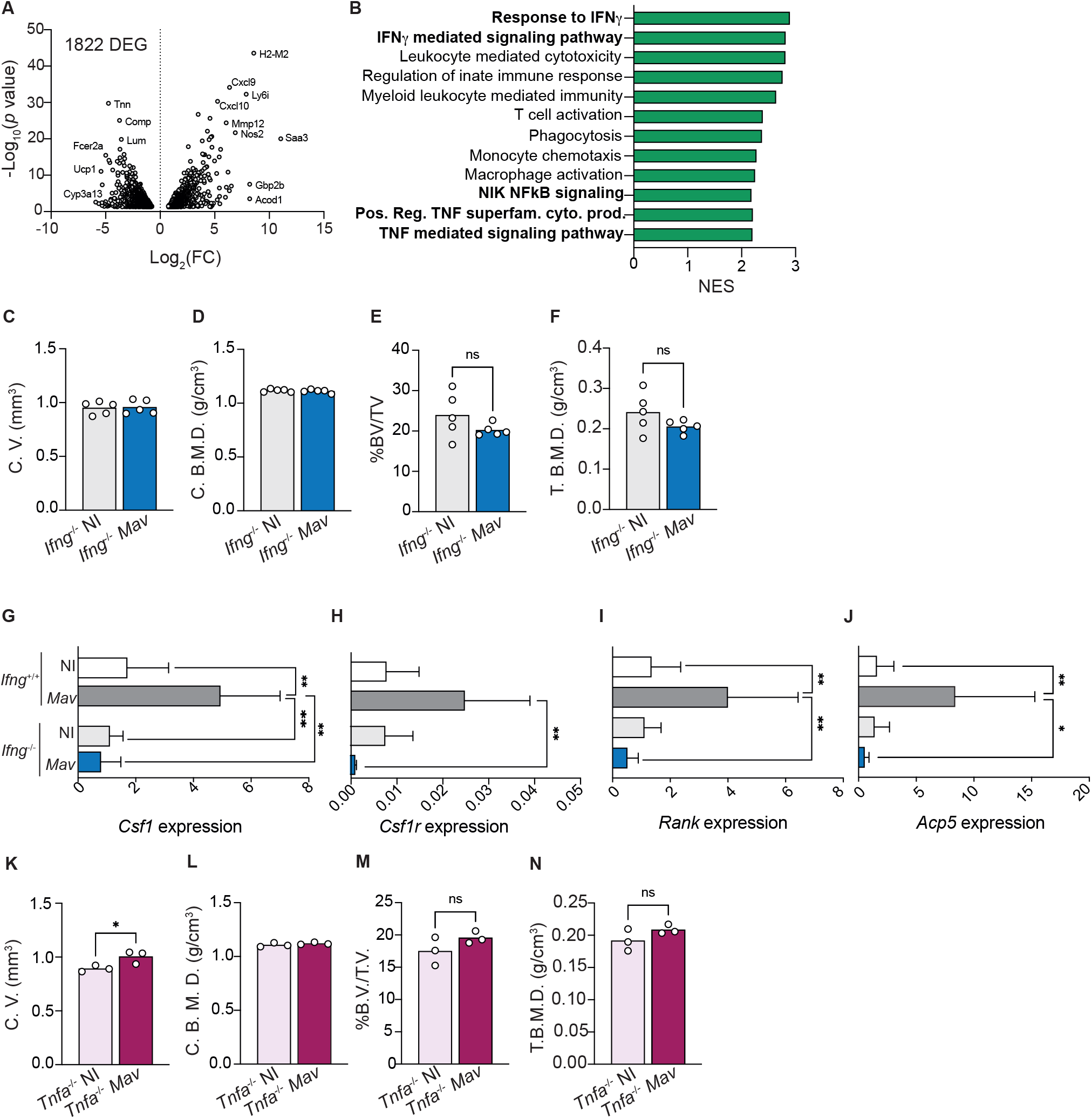
Bone mass reduction during chronic *M. avium* infection is dependent on IFNγ and TNFa production. **A**) Volcano plot of the differentially (and statistically significant) expressed genes in the bone of *M. avium*-infected mice compared to non-infected littermate control mice. **B)** Differentially upregulated pathways (from the GO Biological Process dataset) in the bone of infected mice compared with non-infected controls. **C-D)** Measurement of the cortical volumes (C. V., **C**), and cortical bone mineral density (C. B. M. D., **D**) in *M. avium*-infected (*Mav* bars) compared to non-infected mice (NI bars). **E**-**F**) Measurement of the ratio of bone volume to tissue volume (%BV/TV, **E**), and trabecular mineral bone density (T. B. M. D., **F**). N=5-6 mice per experimental group; data representative of two independent experiments. Bars indicate average; circles depict each individual mouse analyzed. n.s., p>0.0,5 by unpaired Student’s *t* test. **G-J**) Expression of *Cfs1* (**G**), *Csfr1* (**H**), *Rank* (**I**), *Acp5* (**J**), in the bones of infected (*Mav* bars) and non-infected (NI bars) *Ifng^−/−^* and *Ifng^+/+^* mice. Bars represent the average ± standard deviation of the experimental group. *, p<0.0,5, ** p<0.01 by two-way ANOVA. **K-L**) Measurement of the cortical volumes (C. V., **K**), and cortical bone mineral density (C. B. M. D., **L**). **M-N**) Measurement of the ratio of bone volume to tissue volume (%BV/TV, **M**), and trabecular mineral bone density (T. B. M. D., **N**). N=3 mice per experimental group; data representative of two independent experiments. Bars indicate average; circles depict each individual mouse analyzed. n.s., *p*>0.0,5 by unpaired Student’s *t* test.

Given the additional link to TNFa pathways, we decided to investigate the contribution of TNFa to the bone alterations induced by *M. avium* infection. In mice lacking the *Tnfa* gene, *M. avium* infection did not generally result in reduction of cortical and trabecular bone mass and mineral density (**Fig. 2, K** to **N; Fig. S1, Q** to **Z**). Of note, cortical bone mass, trabecular bone volume and trabecular total volume were significantly increased in *Tnfa^−/−^* infected mice (**Fig. 2K**; **Fig. S2**, **X** and **Y**). Together, the data suggest that osteopenia during *M. avium* infection requires both IFNg and TNFa.

### Soluble factors produced by *M. avium-infected* macrophages increase osteoclastogenesis

Next, we questioned whether IFNg and TNFa had a direct effect on osteoclastogenesis. For this purpose, we used two in vitro systems. Firstly, osteoclasts were differentiated in vitro with RANKL and MCSF, the two osteoclastogenesis instructing cytokines, in the absence or presence of IFNg or TNFa. The baseline of differentiation was determined for the condition RANKL and MCSF alone. IFNg addition to in vitro osteoclast cultures did not alter osteoclastogenesis relatively to baseline conditions (**Fig. 3, A** and **B**). Secondly, 3D cultures of osteoclasts using bone slices were performed to determine the bone resorption capacity of the formed osteoclasts. Osteoclasts formed in the presence of IFNg similarly resorbed bone compared to the cultures without IFNg addition (**Fig. 3C**). These data suggest that the role of IFNg in bone loss in vivo is most likely indirect perhaps by the activation of macrophages, leading to the production of soluble factors thus, increasing osteoclastogenesis (Gao et al. 2007, Tang et al. 2018).

**Figure 3.**
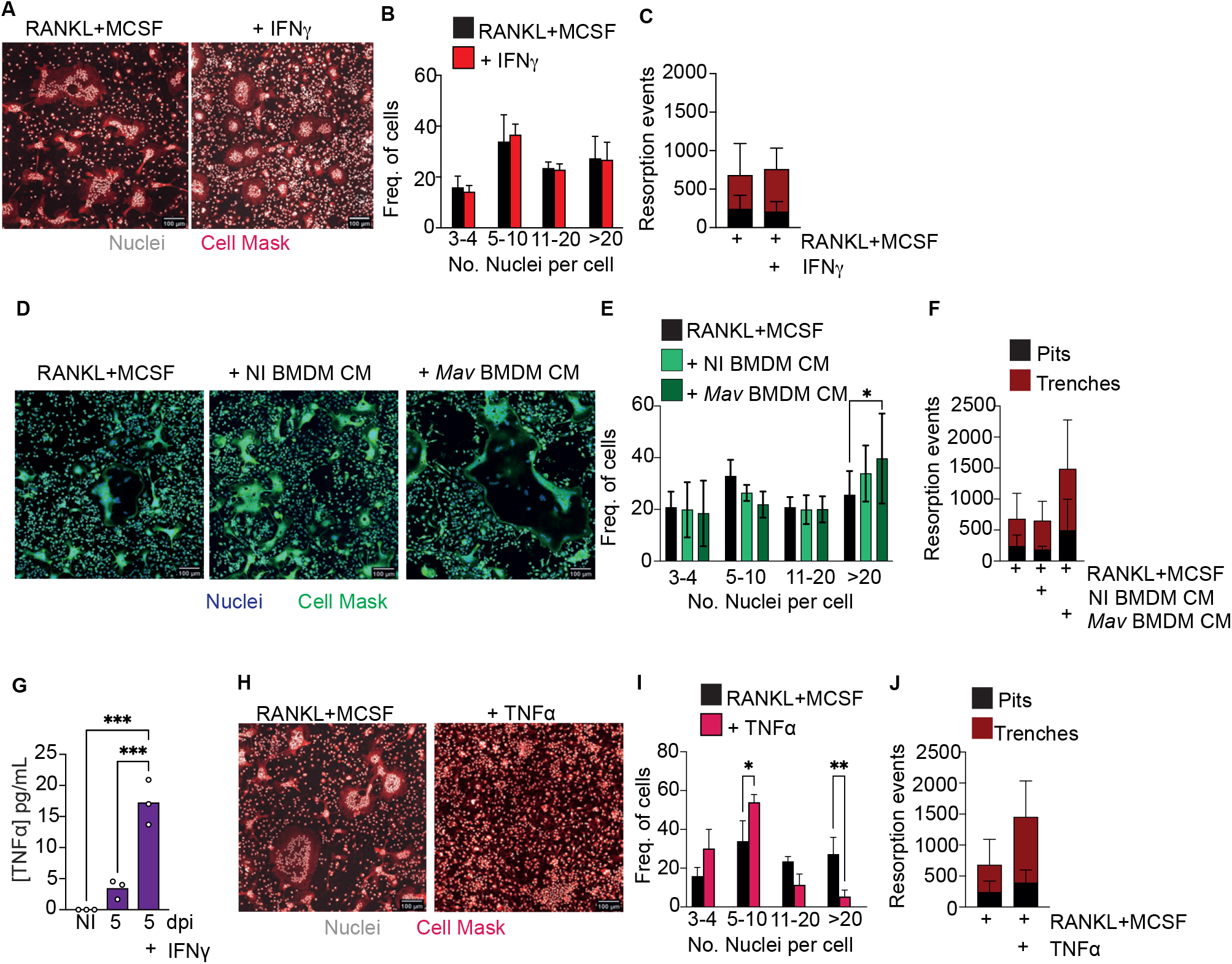
Soluble factors produced by *M. avium-infected* mice increase osteoclastogenesis. **A)** Osteoclast differentiation in the presence of saturating amounts of M-CSF and RANKL (left panel) and with the addition of 1 ng/mL of IFNg. **B**) Frequency of multinucleated cells grouped by number of nuclei per cell. Data are pooled of three independent experiments. Bars represent the average ± standard deviation of the three independent experiments. **C**) Enumeration of pits (black bars) and trenches (red bars). Data are representative of two independent experiments. Bars represent the average ± standard deviation of 4 replicates in each experimental group. **D**)Osteoclast differentiation in the presence of saturating amounts of M-CSF and RANKL (left panel) and with the addition of conditioned media (CM) from non-infected (NI BMDM, middle panel) and *M. avium*-infected bone marrow-derived macrophages (*Mav* BMDM) (right panel). **E**) Frequency of multinucleated cells grouped by number of nuclei per cell. Data are pooled of six independent experiments. Bars represent the average ± standard deviation of the six independent experiments. *, p<0.0,5, by Two-way ANOVA. **F**) Enumeration of pits (black bars) and trenches (red bars). Data are representative of two independent experiments. Bars represent the average ± standard deviation of 4 replicates in each experimental group. **G)** Measurement of TNFa in conditioned media from bone marrow derived and *M. avium* infected-macrophages. **H**) Osteoclast differentiation in the presence of saturating amounts of M-CSF and RANKL (left panel) and with the addition of TNFa (right panel). **I**) Frequency of multinucleated cells grouped by number of nuclei per cell. Data are pooled from four independent experiments. **J**) Enumeration of pits (black bars) and trenches (red bars). Data are representative of two independent experiments. Bars represent the average ± standard deviation of the experimental group. *, p<0.0,5, ** p<0.01 by Two-way ANOVA.

Having seen that IFNg is likely acting through indirect mechanisms, we asked whether infected macrophages in the bone marrow parenchyma could be shaping the immune microenvironment leading to bone loss. So, we next repeated our in vitro experiments supplementing the osteoclast cultures with non-infected or infected BMDM conditioned media. In the presence of conditioned media from infected macrophages, the frequency of osteoclasts with more than 20 nuclei increased (**Fig. 3, D** and **E)**. Moreover, in 3D cultures of osteoclasts on top of bone slices, more pits and trenches were formed in the presence of conditioned media from *M. avium*-infected macrophages (**Fig. 3F**). Together, our data indicate that the local production of cytokines and other soluble factors by macrophages during infection leads to an increased osteoclast production. We next sought to identify these soluble factors.

Taking our above findings showing that both IFNg and TNFa are required as in vivo mediators of bone loss, that IFNg likely acts through indirect mechanisms and that soluble factors produced by infected macrophages induce osteoclast production, we hypothesize that IFNγ may act on macrophages during infection to potentiate TNFa production, which in turn directly or indirectly promotes osteoclast activity. Thus, we firstly measured TNFa production by *M. avium* infected macrophages in the absence or presence of IFNγ. As expected (Appelberg et al. 1994), the production of TNFa was elevated in the presence of IFNγ (**Fig. 3G**). TNFa treatment of osteoclast cultures reduced the number of nuclei per osteoclast (**Fig. 3, H** and **I**), whilst inducing osteoclasts to perform more resorption events, namely trenches (**Fig. 3G**), as compared to baseline conditions. Overall, our results indicate that TNFa directly promote osteoclast activity by increasing the proportion of trenches formed on the bone surface. Contrary to the standard mode of bone resorption (the pit mode), the trenches result from a faster and longer resorption of bone as the osteoclasts move, creating deeper and parallel erosions on the bone surface (Merrild et al. 2015, Soe and Delaisse 2017), suggesting a more aggressive resorption mode. Additionally, our data also indicate that other soluble factors in addition to TNFa and produced by infected macrophages impact osteoclast formation.

### SAA3 is produced during mycobacterial infection impairs new bone formation

In the search for other factors than TNFa involved in increased osteogenesis, we noted that the most differentially upregulated gene in the bones of *M. avium*-infected mice was *Saa3* (**Fig. 4A**). This gene codes for the Serum Amyloid A (SAA)3 protein, which has been recently implicated in bone homeostasis (Thaler et al. 2013, Thaler et al. 2015, Choudhary et al. 2016, Choudhary et al. 2018), namely by enhancing osteoclastogenesis (Thaler et al. 2015). Quantitative real-time PCR analysis confirmed that the expression of *Saa3* is upregulated in the bone during *M. avium* infection (**Fig. 4B**). Moreover, the expression of *Saa3* in the bone was dependent on the production of both IFNγ and TNFa, as mice deficient in either of these cytokines do not upregulate *Saa3* expression in the bone during *M. avium* infection (**Fig. 4, C and D**). Importantly, in mice infected with a low dose of *M. tuberculosis* by the aerosol route, the expression of *Saa3* in the bone marrow was also upregulated as compared with non-infected mice (**Fig. 4E**). These data show that *M. tuberculosis* infection may be paralleling our observations with *M. avium*, and furthermore agree with previous observations that *Saa3* expression is upregulated in bone marrow-derived macrophages and in the lungs after infection with *M. tuberculosis* (Tornack et al. 2017, Mayito et al. 2019).

**Figure 4.**
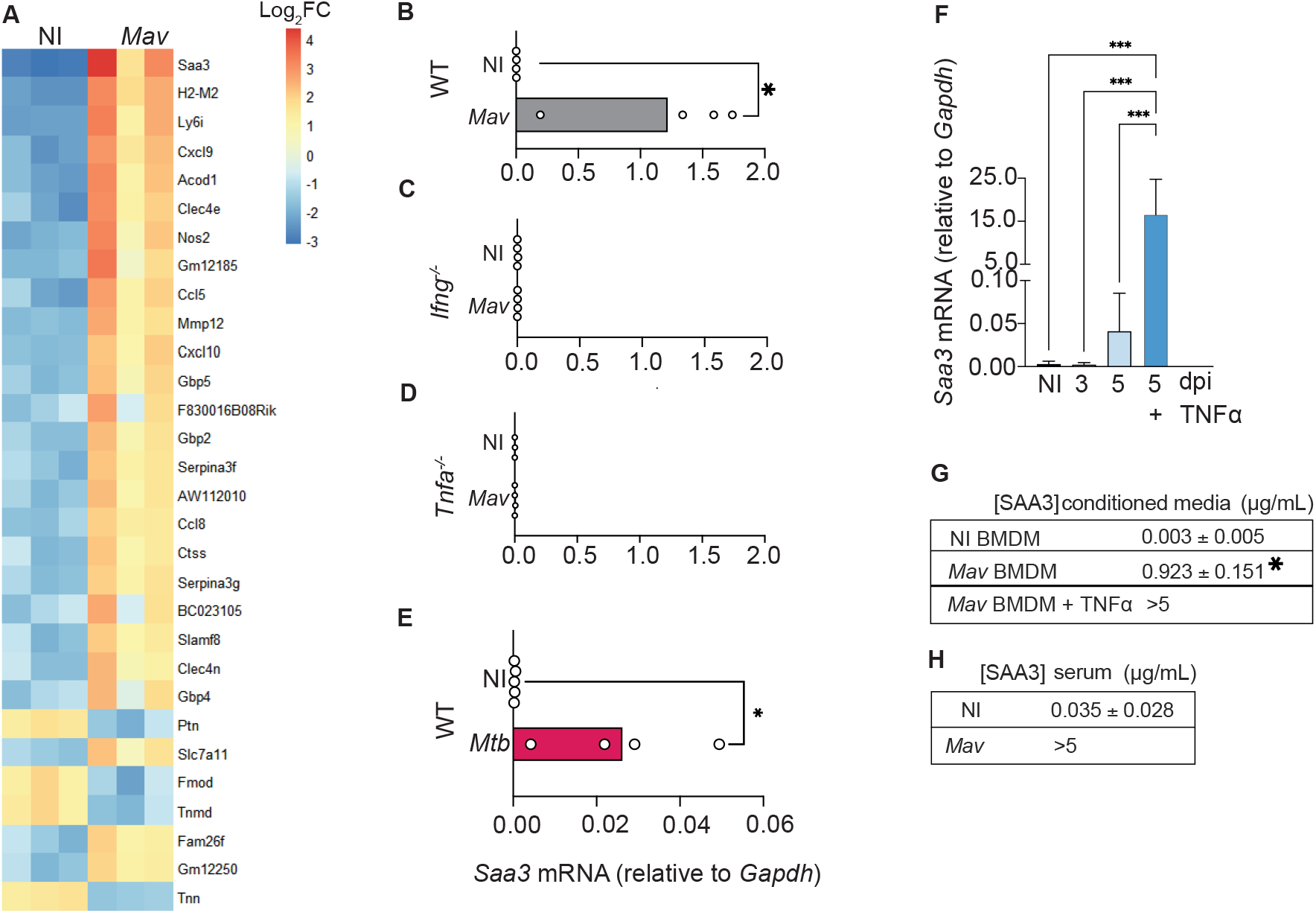
*M. avium*-infected macrophages produce SAA3. **A**) Heatmap representing the 30 most differentially expressed genes in the femurs of *M. avium*-infected mice (*Mav*) compared with non-infected controls (NI). **B-D**) *Saa3* expression relative to *Gapdh* in the femurs of non-infected (white bars) and *M. avium*-infected (gray bars) wild-type (WT, **C**), *Ifng^−/−^* (**D**), and *Tnfa^−/−^* (**E**). Bars indicate mean and circles depict individual mice analyzed. **F**) *Saa3* expression in the bone marrow of mice infected with *M. tuberculosis* by the aerosol route. Bars indicate the average and circles depict each individual mouse. *, p<0.0,5, by Student’s *t* test. **F**) *Saa3* expression relative to *Gapdh* in bone marrow derived macrophages. Bars represent the average ± standard deviation of the 4 replicates in each experimental group. **G**) Quantification of SAA3 in conditioned media from bone marrow derived macrophages, expressed as average ± standard deviation of the experimental group. Results are pooled from 3 independent experiments. *, p<0.0,5, ***, p<0.001 by two-way ANOVA. **H**) Measurement of SAA3 in the serum of *M. avium* - infected mice (*Mav*) compared to non-infected controls (NI) expressed as average ± standard deviation of the experimental group. N= 5 mice per group.

We also found that *Saa3* was upregulated in BMDM 5 days after *M. avium* infection and was further augmented by the addition of TNFa to the macrophages’ cultures (**Fig. 4F**). Corroborating the gene expression data, SAA3 was present in the conditioned media of infected macrophages, and the levels were further increased when infected macrophages were treated with TNFa (**Fig. 4G**), as expected (Betts et al. 1993, Maier et al. 2005, Sommer et al. 2008, Sack 2018).

To determine whether the alterations in *Saa3* expression in the bone correlated with the systemic levels of the cytokine, SAA3 levels were measured in the serum of *M. avium-infected* and non-infected mice. Indeed, high concentrations of SAA3 were found in the serum of infected mice at 8 weeks post-infection (**Fig. 4H)**.

Thus far, our results suggest that during *M. avium* infection, the production of IFNg leads to the activation of macrophages with the concomitant production of TNFa and SAA3. In the absence of TNFa, SAA3 was not expressed in the bone and the bone mass was not altered. Additionally, SAA3 was produced by TNFa-stimulated *M. avium*-infected macrophages.

### SAA proteins are a potential biomarker of bone loss in patients with active Tuberculosis

SAA proteins are well-conserved throughout the mammalian radiation (Uhlar et al. 1994) and have several isoforms (Uhlar and Whitehead 1999). Although the human *Saa3* is a non-translated pseudogene (Kluve-Beckerman et al. 1991), the human SAA1 and SAA2 display a remarkable homology to mouse SAA3 (Thaler et al. 2015). To test if human SAA would also influence osteoclast differentiation, we used our in vitro system incubating osteoclasts in the presence of human recombinant SAA1 and SAA2 proteins (human SAA). Osteoclastogenesis was increased (**Fig 5, A** and **B**) but the number of resorption events was decreased (**Fig. 5C**), indicating that besides SAA other soluble factors are important for the increased bone resorption. According to our mouse data, TNFa is a likely candidate. Indeed, in osteoclasts cultured in the presence of TNFa and human SAA, we not only observed an increase of osteoclastogenesis, but also of the proportion of trenches formed (**Fig. 5, D** and **F**). Overall, these results indicate that SAA and TNFa co-operate as pro-osteoclastogenic factors, which during mycobacteria infection are produced by infected macrophages.

**Figure 5.**
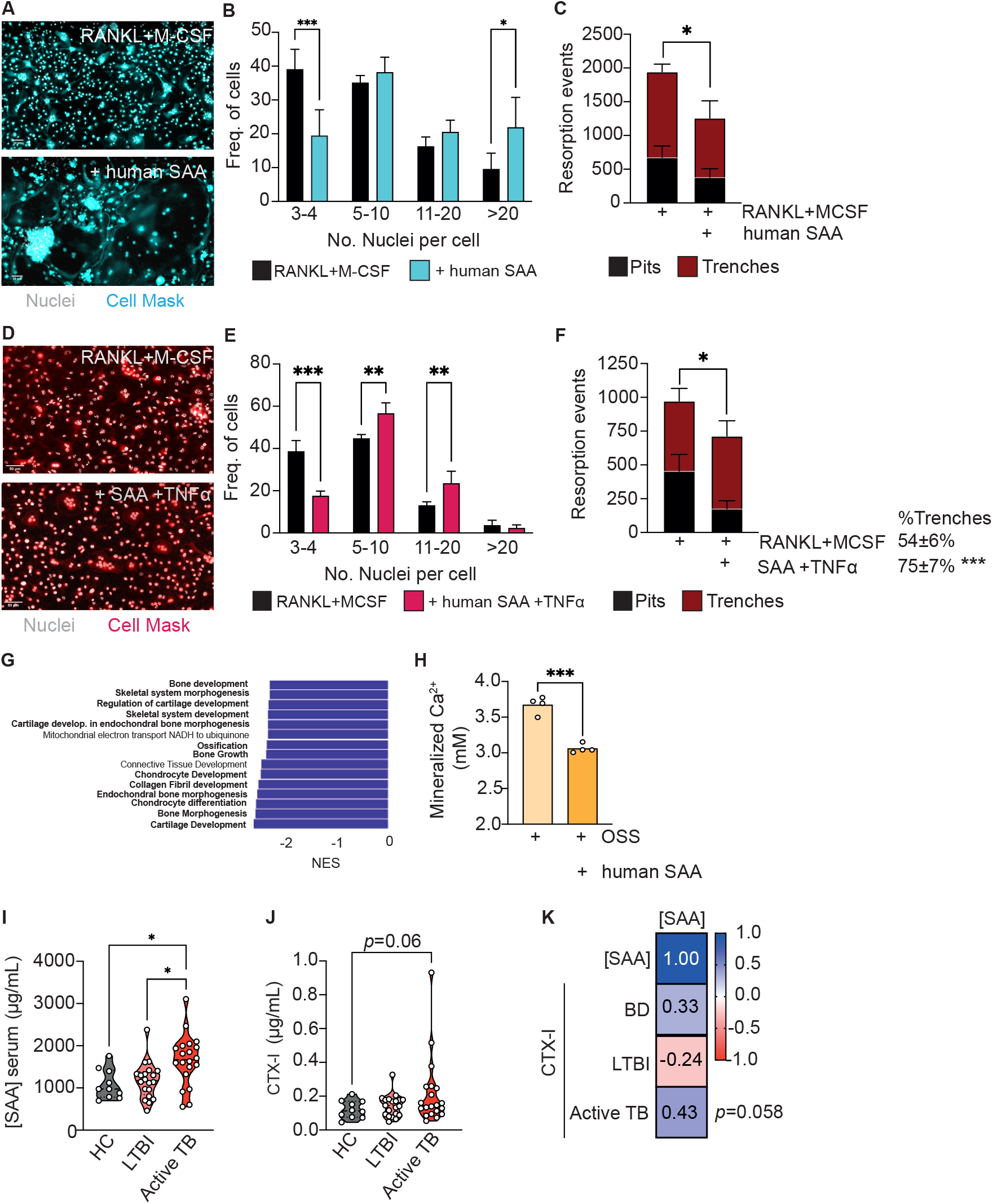
Human SAA increase osteoclastogenesis and decrease osteoblastogenesis and correlate with altered marker of bone turnonver. **A)** Osteoclast differentiation in the presence of saturating amounts of M-CSF and RANKL (top panel) and with the addition of human SAA (bottom panel). Scale bar: 50 μm **B**) Frequency of multinucleated cells grouped by number of nuclei per cell. Data are pooled of three independent experiments. Bars represent the average ± standard deviation of the three independent experiments. **C**) Enumeration of pits (black bars) and trenches (red bars). Data are representative of two independent experiments. Bars represent the average ± standard deviation of 4 replicates in each experimental group. **D)** Osteoclast differentiation in the presence of saturating amounts of M-CSF and RANKL (left panel) and with the addition of human SAA and TNFa (right panel). Scale bar: 50 μm **E**) Frequency of multinucleated cells grouped by number of nuclei per cell. Data are pooled of two independent experiments. Bars represent the average ± standard deviation of the two independent experiments. **F**) Enumeration of pits (black bars) and trenches (red bars). Data are representative of two independent experiments. Bars represent the average ± standard deviation of 4 replicates in each experimental group. *, p<0.0,5, ** p<0.01, ***, p<0.001 by Two-way ANOVA. **G)** Top15 of most differently enriched pathways (from the GO Biological Process dataset) in the bones of infected mice compared to non-infected littermate controls. **H)** Quantification of deposited calcium of murine bone marrow MSC cultured with osteoblast stimulatory supplement (OSS) plus 5 μg/mL human SAA. Bars indicate average; each circle depict each of 4 replicates per experimental group. Experiments are representative of two independent experiments. ***, p<0.001 by Student’s *t* test. **I**) Measurement of SAA in the serum of healthy controls (HC), IGRA+ with latent tuberculosis infection (LTBI) individuals, and patients with active Tuberculosis (active TB). **J**) Measurement of CTX-I in the serum of HC, IGRA-individuals, LTBI patients and patients with active TB. Violin plots depict the density, and each circle represents each patient. *, p<0.0,5, by two-way ANOVA. **D**) Pearson correlations between circulating levels of SAA and CTX-I.

Besides increasing osteoclastogenesis, SAA3 was also reported to inhibit osteoblastogenesis (Thaler et al. 2013, Thaler et al. 2015, Choudhary et al. 2016). Thus, we interrogated our transcriptomic data (**Fig. 2A**) for possible differences in bone morphogenesis, bone development and growth, and ossification pathways and found that they were differentially and statistically downregulated in the femurs of *M. avium*-infected mice compared with the femurs of littermate control mice (**Fig. 5G**). This finding led to the hypothesis that osteoblastogenesis may also be impaired during mycobacterial infections. To test if human SAA might also impact on osteoblastogenesis, MSC from the bone marrow of non-infected mice were isolated and osteoblast formation assays were performed in the presence and absence of human SAA. Osteoblast formation and bone matrix mineralization was detected by alizarin red staining. The addition of human SAA reduced the calcium deposits in the cultures (**Fig. 5H**).

The incidence of non-tuberculous mycobacterial infections, as *M. avium*, has been rising (Donohue and Wymer 2016), namely in cystic fibrosis patients (Viviani et al. 2016) and even in seemingly healthy children and adults (Ratnatunga et al. 2020). However, *M. tuberculosis* remains one of the deadliest human pathogens (WHO 2021). Given the data suggesting a role for SAA in *M. avium* and *M. tuberculosis* mouse infection, as well as for human SAA in modulating osteoclast/osteoblast homeostasis, we went on to investigate whether the levels of SAA might change during active tuberculosis. For that, circulating levels of SAA were measured in the plasma of a cohort of patients with active tuberculosis compared to those detected for healthy controls of unknown IGRA status or for IGRA+ (latent tuberculosis infection, LTBI) individuals. Patients with active tuberculosis showed higher systemic SAA levels as compared to both healthy and LTBI controls (**Fig. 5I**). To correlate the alterations seen in SAA proteins to bone loss, we next measured the circulating levels of CTX-I, as a biomarker of altered bone turnover (Naylor and Eastell 2012), in the plasma of the study cohort. We found an increased amount of CTX-I in active tuberculosis patients which did not reach statistical significance (**Fig. 5J**). Of note, there was a moderate correlation between the systemic levels of SAA and CTX-I which was only observed in the case of active tuberculosis patients (**Fig. 5K**). These findings highlight the potential for the use of the systemic levels of SAA proteins as a biomarker of increased risk of bone loss during chronic mycobacterial infections.

## Discussion

Although it is well established that mycobacteria can infect the bone (Hogan et al. 2019) and that previous mycobacterial infection increases the risk of osteoporosis (Yeh et al. 2016, Choi et al. 2017), very little evidence of the pathophysiology of these alterations has been gathered. Our study shows that during chronic disseminated *M. avium* infection, IFNγ and TNFa induce bone loss, due to both increased resorption and decreased new bone formation. Despite the key role for IFNγ that we show here, it was demonstrated that patients with autosomal dominant IFNγR1 or STAT1 deficiency showed enhanced osteoclastogenesis (Tsumura et al. 2022), which may be explained by the variable penetrance of disease (Gruber and Bogunovic 2020) and by the fact that T cells from these patients still produce TNFa upon stimulation (Kerner et al. 2020), which could mediate the increased osteoclastogenesis. The study of possible bone loss in these patients warrants further research and may shed light into alternative or complementary mechanisms to those reported here. Our work also advances the growing body of evidence supporting a reciprocal interaction between bone homeostasis and hematopoietic development, as the myeloid skewing of hematopoietic stem cells during mycobacterial infection led to an increased osteoclast production. Furthermore, alterations in LEPR+ MSC correlated with decreased bone formation. In line with our observations, a previous study demonstrated that sepsis altered both hematopoietic stem cells and hematopoiesis supporting-MSC leading to a reduction in lymphocyte and osteoblast production (Terashima et al. 2016).

Additionally, we found evidence that SAA proteins strongly associate with bone alterations in both *M. avium* and *M. tuberculosis* infections, being also altered in the serum of TB patients. SAA proteins are described as acute phase proteins that are expressed mostly by hepatic macrophages in the liver (Sack 2018) but also by monocytes and macrophages in other tissues, chondrocytes, adipocytes, and osteoblasts in response to inflammatory stimuli (Meek and Benditt 1986, Meek et al. 1992, Malle and De Beer 1996, Kumon et al. 1997, Yamada et al. 2000, Vallon et al. 2001, Thorn et al. 2003, Zerega et al. 2004, Han et al. 2007, Reigstad et al. 2009). Furthermore, as we show that SAA3 production and bone loss are dependent on inflammatory cytokines, the clinical relevance of SAA proteins as a biomarker of bone loss could be extended to other pathologies such as chronic cardiovascular diseases (Johnson et al. 2004, Katayama et al. 2007), inflammatory rheumatic diseases as well as COVID-19 (Soric Hosman et al. 2020), in which the systemic levels of SAA are elevated. In the long run, these results will also contribute to a deeper understanding of the mechanisms underlying bone loss during inflammatory diseases and consequently to the proposal of new therapeutic strategies to prevent or cure these disorders.

Our results are the first evidence correlating in humans with active tuberculosis the circulating levels of SAA proteins and bone turnover markers. These findings may expand to other settings, where biochemical markers of bone turnover such as CTX-I provide a non-invasive assessment for the clinical assessment and for guiding and monitoring of treatment (Naylor and Eastell 2012). Nevertheless, some studies have reported that despite no alteration in CTX-I levels, alterations in the microstructure and mineral density may be already observed by high resolution peripheral computed tomography (HR-pQCT) and dual X-ray absorptiometry (DXA), respectively (Putman et al. 2014, Gensburger et al. 2016, Mathiesen et al. 2019). As DXA and HR-pQCT are not routinely performed on tuberculosis patients without a previously diagnosed bone disease, the use of CTX-I levels was the available readout of altered bone turnover. Thus, the measurement of the CTX-I in the serum of tuberculosis patients may underestimate the impact of the increased circulating SAA levels in the bone health of these patients. A prospective study to correlate the circulating SAA levels and the bone mineral density in a cohort of tuberculosis patients is warranted.

## Materials and Methods

### Bacteria

*Mycobacterium avium* 25291 SmT (obtained from the American Type Culture Collection, Manassas, VA) was grown at 37°C in Middlebrook 7H9 broth (BD Difco, USA), supplemented with 0.05% of Tween 80 (Sigma, St. Louis, MO) and 10% ADC (Albumin-Dextrose-Catalase). *M. avium* were collected during the exponential growth phase, centrifuged, washed twice with saline containing 0.04% Tween 80, re-suspended in the same solution, and briefly sonicated at low power to disrupt bacteria clumps. *M. tuberculosis* HN878 (clinical isolate) was grown to midlog phase in Middlebrook 7H9 broth supplemented with 0.05% Tween80, and 0.5% glycerol. Aliquots were prepared and stored at −80°C, prior to CFU determination or until needed. Just before use, an aliquot was thawed and diluted to the appropriate concentration. Work with *M. avium* and *M. tuberculosis* was performed under ABSL-2 or 3 facilities, respectively.

### Mice

C57BL/6J, *Ifng^−/−^*, MIIG, and *Tnfa^−/−^* mice were bred and housed under specific pathogen-free conditions at Instituto de Investigação e Inovação em Saúde (i3S) animal facility. Mice were kept inside individually ventilated cages with HEPA filters and fed with sterilized food and water *ad libitum*. Mice were maintained on a 12:12 light cycle at 45-65% humidity and provided ad libitum water and standardized synthetic diet (Envigo Teklad Global Rodent Diet, 2014S). Animal experiments were approved by the i3S Animal Ethics Committee and licensed by Portuguese Competent Authority (DGAV), in July 6th, 2016 with reference 0421/000/000/2016. All animals were handled in strict accordance with good animal practice as defined by national authorities (DGAV, Decreto-Lei 113/2013, August 7th) and European Directive 2010/63/EU. The i3S animal facility is AAALAC accredited and follows the Guide for the Care and Use of Laboratory Animals, principle of the Three R’s, to replace, reduce, and refine animal use for scientific purposes, as well as FELASA recommendations.

### Human samples

The study protocol leading serum isolation was approved by the Portuguese *Comissão de Ética para a Saúde da ARS Norte* (#T792). To ensure confidentiality, each case was anonymized by the assignment of a random identification number. Participants were enrolled at a TB clinic in Porto and provided informed consent. Experiments were conducted according to the principles expressed in the Declaration of Helsinki. 10 healthy controls of unknown IGRA status (6 males and 4 females, 49±7 years old), 20 IGRA+ latent tuberculosis infection (LTBI, 9 males and 11 females, 54±19 years old), and 20 patients with active tuberculosis (9 males and 11 females, 50±17 years old).

### *In vivo* infection

8-to 10-week-old mice were infected with 1 million CFU of *M. avium* 25291 SmT intravenously (iv) into one of the lateral tail veins. Control groups were injected with the same volume of saline solution by the same route. At 8-weeks p.i., mice were anesthetized using an isoflurane chamber. Blood was collected by retro-orbital puncture under deep anesthesia, immediately before euthanasia. Long bones were also collected. For *M. tuberculosis* experiments, mice were infected via the aerosol route using an inhalation exposure system, calibrated to deliver ~200 colony forming units to the lung. The initial bacterial load was determined by quantification of colony forming units in the lungs three days after infection. Infected mice were monitored regularly for signs of illness such as wasting, piloerection and hunching. At 27 days post infection, mice were euthanized by CO_2_ inhalation and long bones were collected. Long bones from aged-matched non-infected mice collected at the same time and used as controls.

### Bone morphometric analysis by microCT

Tibias were harvested, cleaned of muscle, fixed in 4% paraformaldehyde at 4°C, overnight, and scanned at 70Kv, 200 μA with 0.5 mm aluminum filter, and an isotropic resolution of 6 μm, using SkyScan 1276 System (Brucker, Belgium). The obtained projection images were reconstructed with NRecon software, followed by bone alignment along the sagittal axis using DataViewer software. Morphological analysis of both cortical and trabecular bone were performed with CTAnalyser software, following the recommended guidelines (Bouxsein et al. 2010). Cortical analysis were carried out in 150 slices at the diaphysis (700 slices away from distal growth plate). 3D images of representative samples were generated using CTVol software.

### 2D Histomorphometry analysis

Tibias/femurs were cleaned from surrounding soft tissue and fixed in 10% (v/v) PFA for 24 h and rinsed in phosphate buffered saline (PBS) solution at 4°C. Samples were then dehydrated in serial ethanol solutions (50–100%) for 3 days each, cleared in xylol for 24 h and further embedded in methyl methacrylate. Undecalcified 7 μm sagittal cuts were stained with modified Masson-Goldner Trichrome, and processed for static histomorphometric analysis as described elsewhere(Sousa et al. 2012), using OsteoMeasure software (OsteoMetrics, Decatur, GA, USA). All histomorpometric analysis were performed by a single blinded operator. Digital images were obtained using NanoZoomer 2.0HT (Hamamatsu) and representative images were acquired.

### Dynamic in vivo bone labeling

Double labeling with xylenol orange and calcein was used to study bone turnover during *M. avium* infection(van Gaalen et al. 2010). Thus, 6 weeks post-infection, a saline solution of xylenol orange (Sigma-Aldrich, USA) was subcutaneously administered at a dose of 80 mg/kg. Two weeks later, a saline solution of calcein green (Sigma-Aldrich, USA) at a dose of 15 mg/kg was administered by the same route as xylenol orange. Three days later (at 8 weeks post-infection), mice were euthanised, and femurs and tibias were collected, fixed in 4% paraformaldehyde (BioOptica, Italy) at 4°C, overnight. Then, the bones were washed 3 times with PBS1x for 10 min, followed by dehydration with PBS1x/ 30% sucrose, at 4°C, overnight. Long bones were embedded in OCT (ThermoFisher, UK), snap-frozen in an ethanol/dry ice bath, and 20-μm sections were cut using the Kawamoto method(Kawamoto and Kawamoto 2021), using a Leica CM 3050S cryostat (Leica

Biosystems, Portugal). Sections were kept at −20°C, inside a silica gel chamber, for at least 12 hours. Then, slices were warmed to room temperature inside a silica gel chamber and mounted in a 30% glycerin solution. Images were acquired on a Leica SP8 confocal microscope (Leica Microsystems, Germany), using LASX software (Leica Microsystems, Germany). Images were analyzed using the measurement tool of Imaris Software (Oxford Instruments, UK).

### Immunofluorescence of femur whole mounts

Long bones were collected from infected mice at 8 weeks post-infection and from littermate non-infected controls, fixed, dehydrated, and embedded in OCT, as described before. Then, bones were trimmed in a cryostat to expose the medullary cavity, and OCT was removed by melting at room temperature. After 3 washes with PBS for 10 minutes, to block the non-specific binding and to permeabilize, whole mounts were incubated for 1 hour at room temperature with PBS containing 20% fetal bovine serum and 0,5% Triton X-100 (Sigma Aldrich, USA). Afterwards, whole mounts were stained for 3 days, at room temperature, with rabbit anti-TRAP antibody (1:50 dilution, Abcam, UK). Whole mounts were washed in PBS for 10 minutes, three-times, and stained with Alexa 647 anti-rabbit antibody (1: 500 dilution, Jackson ImmunoResearch, UK) and DAPI (Sigma-Aldrich) for 3 days, at room temperature and protected from the light. After 3 washes in PBS for 10 minutes, whole mounts were transferred to an imaging chamber (ibidi GmbH, Germany) containing 30% glycerin solution and imaged on a Leica SP8 confocal microscope (Leica Microsystems, Germany). Bone surface expressing TRAP was quantified using Imaris Software (Oxford Instruments, UK).

### Histology

Femurs were fixed in 10% neutral buffered formalin solution overnight at 4°C, in EDTA/Glycerol solution for 3 weeks at 4°C, and then included in paraffin blocks. 5-μm thick sections were cut, deparaffinized and processed through downgraded alcohols, and rehydrated.

#### Immunohistochemistry of F4/80 and Ziehl-Neelsen stain

Sections were deparaffinized and processed as described before. To unmask the antigenic epitope, sections were incubated in citrate buffer pH 6.0 (Thermo Fisher Scientific) in a steamer for 20 min, allowed to cool for 10 min at RT, followed by enzymatic digestion with 0.05% Trypsin (Gibco) in a humidified chamber at 37°C for 30 min. Enzymatic reaction was stopped with incubations in cold water for 5 minutes, twice. After blocking of endogenous peroxidase activity using Ultravision hydrogen peroxide block (Thermo Fisher Scientific), endogenous biotin was blocked (Lab Vision Avidin biotin Blocking Solution, Thermo Fisher Scientific). Unspecific binding sites were blocked with Normal Goat Serum (1:5 dilution, DAKO) in antibody diluent (Thermo Fisher Scientific) at RT for 30 minutes. Rat anti-mouse F4/80 antibody (clone BM8, Biolegend) was added to the sections and incubated in a humidified chamber overnight at 4°C. Then, sections were stained with a biotinylated goat anti-rat IgG (Enzo) in a humidified chamber at room temperature for 30 minutes, followed by incubation with Streptavidin-HRP (Vector Laboratories) in a humidified chamber at RT for 30 minutes. DAB Quanto (Thermo Fisher Scientific) was added to allow the color development for 3 minutes. Slides were stained with Carbol Fuchsin and then heated and rinsed off in tap water. Then, 1% of hydrochloric acid in methanol was added to slides. Slides were counter stained with Hematoxylin, rinsed twice with water, dehydrated (reverse of downgraded alchools and deparaffinization), and coverslipped in Entellan. Sections were scanned in D-sight Plus f2.0 and analyzed with D-Sight viewer.

### Flow cytometry

Cells were isolated from bone marrow of long bones by flushing in DMEM supplemented with 2% heat-inactivated bovine fetal serum, 5% HEPES, and PenStrep (all from Invitrogen). Cells were counted using the trypan blue (Sigma-Aldrich) exclusion assay.

#### Myeloid cell analysis

One million BM cells were stained with anti-Gr-1 (clone RB6-8C5, BioLegend), anti-CD115 (clone AFS98, BioLegend), anti-F4/80 (clone BM8, BioLegend) and anti-CD169 (clone 3D6.112, BioLegend). for 30 min, on ice. Cells were washed twice at 2000 rpm for 2 min and fixed with 2% PFA for 10 min at room temperature. Cells were gated as previously described(Chow et al. 2011). Neutrophils: Gr-1^hi^ CD115^−^; Monocytes: Gr1^lo^CD115^+^ F4/80^+^; Macrophages: Gr1^lo^ CD115^−^ F4/80^+^ CD169^+^.

#### Stromal cells analysis

The isolation of bone marrow stromal cells was performed as described before (Cordeiro Gomes et al. 2016). Briefly, long bones were carefully flushed using HBSS supplemented with 2% heat-inactivated fetal bovine serum, PenStrep, HEPES and 200 U/mL Collagenase IV (Sigma-Aldrich). The flushed bone marrow was then digested at 37°C for 20 minutes. Cells clumps were dissociated by gentle pipetting, followed by a further incubation for 10 min at 37°C. Cells were then washed with HBSS/2%FBS, spun at 1000 rpm for 7 min and stained. LEPR staining was performed using biotin-conjugated anti-LEPR (R&D systems) for 1 hour, on ice. The remaining staining using PE-conjugated anti-CD45 (clone 30-F11, BioLegend) and anti-TER119 (BioLegend) and APC-conjugated anti-CD31 (clone 391, BioLegend) and anti-CD144 (BV13, BioLegend) were done for 20 min, on ice, followed by 2 washes at 1000 rpm for 7 minutes. Finally, cells were washed and fixed with 2% PFA for 10 min at room temperature. Cells were analyzed using a BD FACS Canto II flow cytometer and FlowJo.

### RNA isolation and Quantitative Real Time PCR

For *M. avium*-infected mice and littermate non-infected mice, bones were harvested and snap frozen in liquid nitrogen. Bones were crushed in a liquid nitrogen bath and total RNA was isolated using RNA Mini Kit (Invitrogen), according to manufacturer instructions. For *M. tuberculosis*-infected mice bone marrow was flushed from femurs and tibias and extracted with Trizol. cDNA was then synthetized using NZY First-Strand cDNA synthesis kit (NZYTech, Portugal).

Primers sequences:

*Acp5* Fw: 5’-GCTGGAAACCATCATCACCT-3’;

*Acp5* Rv: 5’-TGAAGCGCAAACGGTAGT-3’;

*Csf1* Fw: 5’-GGGCCTCCTGTTCTACAAGT-3’;

*Csf1* Rv: 5’-AGGAGAGGGTAGTGGTGGAT-3’;

*Csf1r* Fw: 5’-TTGGACTGGCTAGGGACATC-3’;

*Csf1r* Rv: 5’-GGTTCAGACCAAGCGAGAAG-3’;

*Gapdh* Fw: 5’-TGTGTCCGTCGTGGATCTGA-3’;

*Gapdh* Rv: 5’-CCTGCTTCACCACCTTCTTGA-3’;

*Rank* Fw: 5’-GTGCTGCTGGTTCCACTG-3’;

*Rank* Rv: 5’-CCGTCCGAGATGCTCATAAT-3’;

*Saa3* Fw: 5’-ACATGTGGCGAGCCTACTCT-3’;

*Saa3* Rv: 5’-GAGTCCTCTGCTCCATGTCC-3’.

### Targeted Transcriptomic Analysis

Whole bone total RNA was isolated from C57BL/6J mice at 8 weeks post-infection and from littermate non-infected mice, as described above. The transcriptomic laboratorial processing was performed by the Genomics Core Facility at i3S (Porto, Portugal). Briefly, RNA concentration was measured using Qubit3.0 fluorometer and total RNA integrity number was determined using the Agilent 2100 Bioanalyzer. Libraries were constructed according to Ion AmpliSeq™ Transcriptome Mouse Gene Expression Kit protocol (Thermo Fisher Scientific, USA) and pooled with each sample ligated to a unique barcode. The pooled libraries were processed on Ion Chef™ System and the resulting 540™ chip was sequenced on Ion S5™ XL System. FASTQ and/or BAM files were generated using the Torrent Suit plugin FileExporter v5.12.

The statistical analysis and graphical representations were performed in Rstudio version 1.4.1106. The quality control of the expression profiles of triplicates of each experimental group were investigated through Principal Component Analysis (PCA). Differentially expressed genes in the bone between infected and non-infected bones were identified by the R package DESeq2 package v1.24 with default parameters (Love et al. 2014), considering an adjusted p-value threshold below 0.05 and a log2FoldChange>0.5. Clustering and heatmap representation of these significantly expressed genes, between uninfected and infected tests, were obtained using the R package heatmap version 1.0.12 package. Pre-ranked pairwise gene-set enrichment analyses (GSEA) were conducted in GSEA software for the Gene Ontology (GO) (biological process or BP, cellular component or CC, and molecular function or MF) and KEGG pathway datasets, after converting the mouse genes in the homologous human genes by using information from the Mouse Genome Informatics Web Site (Bult et al. 2019). The resulting enriched pathways were ordered by the normalized enrichment score (NES) and the false discovery rate (FDR) q-value. Top significantly enriched pathways (FDR < 0.05) were further explored based on information contained in the publicly available database of Ingenuity (https://targetexplorer.ingenuity.com/index.htm; Qiagen, Hilden, Germany).

### Measurement of SAA, TNFa and CTX-I by ELISA

Blood was collected from the orbital vein to non-heparinized tubes and let to clot and spun at high speed to collect serum. Serum levels of SAA3, TNFa and CTX-I were measured using the mouse SAA-3 ELISA kit (Sigma-Aldrich), LEGEND MAX™ Mouse TNF-α ELISA Kit (BioLegend) and the RatLapsTM EIA CTX-I (Immunodiagnostic Systems, Denmark), following the manufacturer’s instruction.

Human peripheral blood was collected by venous puncture and plasma was recovered. SAA and CTX-I were measured using the human SAA ELISA (Sigma-Aldrich) and Serum Crosslaps (CTX-I) ELISA (ImmunoDiagnosticSystems, Denmark), following the manufacturer’s instruction.

### Differentiation of Bone Marrow-derived Macrophages

Bone marrow was harvested from the long bones of C57BL/6 mice by flushing with Hank’s Balanced Salt Solution (HBSS, Gibco, UK). Cells were cultured overnight in DMEM containing 10% fetal bovine serum, 1% pyruvate, 1%-glutamine, 1% HEPES, and 10% L929 cell-conditioned medium (LCCM), at 37°C/ 5% CO_2_. Non-adherent cells were collected and then plated at a cellular density of 5×10^5^ cells/mL. Cells were cultured in DMEM containing 10% fetal bovine serum, 1% pyruvate, 1%-glutamine, 1% HEPES, and 10% L929 cell-conditioned medium (LCCM), at 37°C/5% CO_2_. At day 4 of culture, 10% LCCM was added and at day 7, the medium was renewed. At the 10^th^ day, cell culture media was collected to obtain non-infected bone marrow-derived macrophage conditioned media (NI BMDM CM). Then, cells were in vitro infected with 5×10^5^ CFU/mL of *M. avium* 25291 SmT, for 4 hours at 37°C/5% CO_2_. Afterwards, cells were washed 4 times with warm HBSS and then incubated at 37°C/5% CO_2_ for 5 days. For TNFa and IFNg treatments, a concentration of 10 ng/mL and 16ng/mL, respectively, was added to the cultures for 3 days post-infection. At day 5 post-infection, cell culture medium was removed and filtered to obtain *M. avium*-infected bone marrow-derived macrophages conditioned medium (*Mav* BMDM CM). Conditioned media were stored at −80°C until used. RNA was harvested from infected and non-infected cultures using RNA Mini Kit (Invitrogen). Gene expression was determined as previously described.

### Osteoclast differentiation assays

Hematopoietic progenitors were harvested from the bone marrow by flushing the long bones with aMEM supplemented with 10% fetal bovine serum. After red blood cell lysis, cells were cultured overnight in aMEM supplemented with 10% fetal bovine serum and 10 ng/mL M-CSF, at 37°C/5% CO_2_. Non-adherent cells were collected and then cultured at a cell density of 7.5×10^5^ cells/mL, with 100ng/mL RANKL (BioLegend), 100 ng/mL M-CSF (BioLegend), and 50% NI or *Mav* BMDM CM. Media was renewed every other day. At the 7^th^ day of culture, cells were fixed with 4% paraformaldehyde (BioOptica, Italy) for 10 min, at room temperature, and permeabilized with 0.1% Triton X-100 in PBS 1x for 15 minutes at room temperature. Nuclei were labelled using DAPI, and cytoplasm was labeled with HCS CellMask (Invitrogen, USA). Imaging was performed using IN CELL Analyzer 2000 microscope (GE Healthcare, USA). Osteoclasts were manually quantified using Image J and Microsoft Excel (Microsoft Corporation, USA). An osteoclast was defined as a multinucleated cell with more than 3 nuclei per cell. Four ranges were defined: 3-4 nuclei per cell, 5-10 nuclei per cell, 11-20 nuclei per cell and more than 20 nuclei per cell. The frequency of each class was calculated. To determine osteoclast resorptive capacity, osteoclast precursors were seeded on top of commercially available bovine cortical bone slices (boneslices.com, Denmark). At day 7, cells were lysed by incubation with water. Bone slices were washed and stained with Toluidine Blue. The total number of pits and trenches per slice was enumerated using an optical microscope.

### Osteoblast cultures

Bone marrow stromal cells were flushed from the long bones C57BL6 mice with aMEM supplemented with 10% fetal bovine serum, plated in 75 cm2 flasks and cultured for 7 days. Medium was renewed every 2-3 days to remove hematopoietic cells in suspension. Primary osteoprogenitor cells were harvested at pre-confluence using trypsin solution and cell viability was determined by the Trypan Blue Exclusion assay. Then, cells were plated at a 5×10^5^ cells/mL, under osteogenic conditions (MesenCult Osteogenic Stimulatory Kit, Stem Cell Technologies, USA). Medium was renewed every 3 days. At day 12 of culture under osteogenic conditions, cells were fixed with 4% paraformaldehyde for 10 min, at room temperature, and stained with alizarin-red staining solution (Sigma-Aldrich, USA) for 30 min, at room temperature with gentle shaking. Then, calcium deposits were eluted with 10% cetypyridinium chloride in sodium phosphate (pH 7.0) and quantified at 570 nm.

### Statistics

Student’s two-tailed t test and One-Way ANOVA were performed using GraphPad Prism 9 (GraphPad Software, CA, USA). A p value <0.05 was considered significant.

## Data availability

The RNAseq data is available in GEO under the accession number GSE215856 and can be accessed using the following token: edwpcwserhgxhuf.

## Acknowledgements

This article is a result of the project HEALTH-UNORTE: Setting-up biobanks and regenerative medicine strategies to boost research in cardiovascular, musculoskeletal, neurological, oncological, immunological and infectious diseases (NORTE-01-0145-FEDER-000039), supported by Norte Portugal Regional Operational Programme (NORTE 2020), under the PORTUGAL 2020 Partnership Agreement, through the European Regional Development Fund (ERDF). This work was supported by KOG-202108-00929 from the European Haematology Society, awarded to ACG. Work in the MS lab was financed by FEDER - Fundo Europeu de Desenvolvimento Regional funds through the COMPETE 2020 - Operacional Programme for Competitiveness and Internationalisation (POCI), Portugal 2020, and by Portuguese funds through FCT - Fundação para a Ciência e a Tecnologia/Ministério da Ciência, Tecnologia e Ensino Superior in the framework of the project POCI-01-0145-FEDER-028955 (PTDC/SAU-INF/28955/2017).

The authors acknowledge the support of the i3S Scientific Platform BioSciences Screening, member of the PT-OPENSCREEN (NORTE-01-0145-FEDER-085468) and PPBI (PPBI-POCI-01-0145-FEDER-022122) as well as the support of the i3S Scientific Platform Bioimaging, member of the national infrastructure PPBI - Portuguese Platform of Bioimaging (PPBI-POCI-01-0145-FEDER-022122), for microCT imaging acquisition. The authors also thank to the valuable collaboration of the following i3S Scientific Platforms: Cell Culture and Genotyping Core Facility, Advance Light Microscopy, Genomics, Histology and Electron Microscopy, Animal Facility, and Flow Cytometry Unit. Mariana Resende and Rui Appelberg kindly provided the *Tnfa* deficient mice. The authors thank Dr. João P. Pereira (Yale University) for the critical reading of the manuscript.

ACG and MS are supported by an Individual Scientific Employment contract (CEECIND/00048/2017; CEECIND/00241/2017 respectively). DMS acknowledges the Portuguese Foundation for Science and Technology (FCT) for the Post-Doc fellowship (SFRH/BPD/115341/2016). RJP, DS and AIF have PhD grants (SFRH/BD/145217/2019; SFRH/BD/143536/2019; 2020.05949.BD, respectively) financed by FCT.

Schemes and graphical abstract were created with BioRender.com.

## Author contributions

ACG, ML and MSG supervised the study. ACG conceptualized, designed and performed the experiments. ACG, MS and MSG wrote the manuscript. DMS conceptualized and performed some experiments. TCO, OF, DS, ACM and TS performed experiments. AF, MJT and MS provided human samples. RP and LP performed the transcriptomic analysis.

---

**Figure S1. *Mycobacterium avium* infection increases bone degradation and reduces bone formation. A)** Representative scheme of in vivo *M. avium* infection. **B-C)** Representative image of the cortical (**B**) and trabecular (**C**) tibial bone of *M. avium*-infected (*Mav* bars) and non-infected mice (NI bars) scanned by micro-CT. **D)** Quantification of CTX-I in serum. **E)** Enumeration of the osteoblast number per bone perimeter (N.Ob./B.Pm.) in infected mice (*Mav* bars) *vs* non-infected controls (NI bars). **F)** Enumeration of the frequency of LEPR+ MSC in the bone marrow of infected and non-infected mice by flow cytometry. N=5-6 mice per experimental group; data representative of two independent experiments. Bars indicate average and each dot represents each mouse. **G**) Representative image of the cortical tibial bone of *M. avium-infected (Mav* bars) and non-infected (NI bars) *Ifng^−/−^* mice scanned by micro-CT. **H-L)** Measurement of the marrow volume (M. V., **A**), cortical thickness (**I**), endocortical perimeter (Endo. P., **J**), periosteal perimeter (Perio. P., **K**), and mean polar moment of inertia (M. P. M. I., **L**) in *M. avium*-infected *Infgamma^−/−^* mice (*Mav* bars) compared to non-infected littermate controls (NI bars). **M)** Representative image of the trabecular tibial bone of *M. avium*-infected (*Mav* bars) and non-infected (NI bars) *Ifng^−/−^* mice scanned by micro-CT. **N-P**) Measurement of the trabecular bone volume (T. B. V., **N**), trabecular tissue volume (T. T. V., **N**), trabecular number (#T, **P**) in *M. avium*-infected *Infg^−/−^* mice (*Mav* bars) and non-infected controls (NI bars). **Q)** Representative image of the cortical tibial bone of infected and non-infected *Tnfa^−/−^* mice scanned by micro-CT. **R-V)** Measurement of the marrow volume (M. V., **R**), cortical thickness (**S**), endocortical perimeter (Endo. P., **T**), periosteal perimeter (Perio. P., **U**), and mean polar moment of inertia (M. P. M. I., **V**) in *M. avium*-infected *Tnfa^−/−^* mice (*Mav* bars) compared to non-infected littermate controls (NI bars). **W**) Representative image of the trabecular tibial bone of infected and non-infected *Tnfa^−/−^* mice scanned by micro-CT. **X-Z**) Measurement of the trabecular bone volume (T. B.V., **X**), trabecular tissue volume (T.T.V., **Y**), trabecular number (#T, **Z**) in *M. avium*-infected *Tnfa^−/−^* mice (*Mav* bars) and non-infected controls (NI bars). N=3-6 mice per experimental group; data representative of two independent experiments. Bars indicate average; each dot depicts each individual mouse analyzed **i**\*, p<0.0,5, ** p<0.01, ***, p<0.001 by Student’s *t* test.

**Figure S2. In infected bones, *M. avium* resides mostly inside macrophages distributed throughout the bone marrow parenchyma.** 5-μm thick sections of non-infected (NI, left panel) and infected (*Mav*, right panel) femurs stained with anti-F4/80 and Ziehl-Neelsen. (#) mark F4/80+ (brown) cells infected with *M. avium* (red rods).

